# Nanocarrier Drug Release and Blood-Brain Barrier Penetration at Post-Stroke Microthrombi Monitored by Real-Time Förster Resonance Energy Transfer-Based Detection System (FedEcs)

**DOI:** 10.1101/2024.02.28.582471

**Authors:** Igor Khalin, Nagappanpillai Adarsh, Martina Schifferer, Antonia Wehn, Valeria J. Boide-Trujillo, Uta Mamrak, Joshua Shrouder, Thomas Misgeld, Severin Filser, Andrey Klymchenko, Nikolaus Plesnila

## Abstract

Nanotechnology holds great promise for improving the delivery of therapeutics to the brain. However, current approaches often operate at the organ or tissue level and are limited by the lack of tools to dynamically monitor cargo delivery *in vivo*. We have developed highly fluorescent lipid nanodroplets (LNDs) that enable tracking of nanocarrier behaviour at the subcellular level while also carrying a Förster resonance energy transfer (FRET)-based drug delivery detection system (FedEcs) capable of monitoring cargo release *in vivo*. Using two-photon microscopy, we demonstrate that circulating LNDs in naïve mouse brain vasculature exhibit 3D real-time FRET changes, showing size-dependent stability over two hours in blood circulation. Further, in a novel nano-stroke model, dynamic intravital two-photon imaging revealed that LNDs accumulated within cerebral post-ischemic microthrombi, where they released their cargo significantly faster than in normal blood circulation. Furthermore, the blood-brain barrier (BBB) became permeable at the microclot sites thereby allowing accumulated FedEcs-LNDs to cross the BBB and deliver their cargo to the brain parenchyma. This microthrombi-associated translocation was confirmed at the ultrastructural level via volume correlative light-electron microscopy. Consequently, our FedEcs represents a novel tool to quantitatively study the biodistribution and cargo release of nanocarriers at high resolution in real time. By enabling us to resolve passive targeting mechanisms post-stroke, - specifically, accumulation, degradation and extravasation via post-stroke microthrombi - this system could significantly enhance the translational validation of nanocarriers for future treatments of brain diseases.

## Introduction

Brain diseases, including stroke, brain trauma, neurodegenerative disorders, and brain tumors, pose significant global health challenges due to their high morbidity and mortality rates^1^. Effective treatment of these conditions is often hindered by the blood-brain barrier (BBB), a highly selective barrier that restricts the entry of most therapeutics into the brain parenchyma^2^. Overcoming the BBB to deliver drugs effectively remains a critical hurdle in neuroscience and pharmacology.

Nanotechnology is currently revolutionizing the delivery of therapeutics by offering innovative solutions to transport drugs to the target organs. Along with widely used lipid nanoparticles (LNPs), lipid nanodroplets (LNDs), also known as nano-emulsions, are particularly attractive and promising platforms because of their biocompatibility, structural similarity to low-density lipoproteins and favorable pharmacokinetics properties^3^. The liquid core serves as an excellent reservoir for encapsulation of lipophilic drugs and contrast agents^3^, while their lipid shell is biocompatible and therefore generally recognized as safe (GRAS) for the application in patients^4-6^. LNDs are characterized by prolonged circulation time in blood, which enhance their potential for both passive and active targeting tissues^7^. Previous study have demonstrated that LNDs can remain intact in blood circulation for several hours post-injection and preferentially accumulate in tumors due to the enhanced permeability and retention (EPR) effect^7^. However, despite these advantageous properties, investigating the biodistribution of LNDs and other nanoparticles (NPs) in the brain remains a fundamental challenge. Current detection modalities lack the resolution and specificity to dynamically monitor whether NPs cross the BBB, enter the brain parenchyma, and release their therapeutic cargo at the specific site^2^. To overcome this limitation, we developed highly fluorescent LNDs capable to be detected *in vivo* with high spatial resolution. Moreover, our LNDs carry a <u>F</u>örster-resonance <u>e</u>nergy transfer (FRET)-based drug delivery <u>d</u>ete<u>c</u>tion <u>s</u>ystem (FedEcs) which allows for real-time visualization of cargo release *in vivo*. Additionally, the lipid core of LNDs made them visible in electron microscopy (EM), enabling us to establish a novel correlative light-electron microscopy (CLEM) method. This method allows us to combine dynamic changes observed intravitally with ultrastructure insights of the BBB leakage.

Using advanced imaging techniques such as *in vivo* two-photon microscopy and volume CLEM, we investigated the biodistribution and cargo release of FedEcs-LNDs in a mouse model of cerebral ischemia induced by magnetic nanoparticles (a novel nano-stroke model). Our observations revealed that 30 nm FedEcs-LNDs accumulate within microthrombi forming in brain capillaries post-stroke, cross the compromised BBB at these sites, and release their cargo both within the microthrombi and in the brain parenchyma. These findings demonstrate the potential of LNDs for passive targeting of ischemic brain, crossing BBB via microthrombi and highlight the importance of subcellular-level real-time monitoring for the development of effective nanomedicines.

## Results

### Development of highly fluorescent lipid nanodroplets (LNDs) carrying a Förster-resonance energy transfer (FRET)-based cargo delivery detection system (FedEcs)

We aimed to design highly fluorescent nanodroplets which employ FRET to dynamically assess both the structural integrity of the particles and the release of their encapsulated cargo. To achieve this, LNDs were formulated using LabrafacWL and Cremophor ELP® through a spontaneous nanoemulsification^8^. These LNDs were loaded with two fluorophores capable undergo FRET and detect the LNDs decomposition: upon degradation of the LNDs and/or cargo release, the distance between the donor and acceptor dyes increases thereby leading to the loss of FRET and a subsequent reduction of the ratio between donor and acceptor fluorescence intensity **(Figure 1A)**. The fluorescent dye F888, which was specially designed for high loading into LNDs and for superior fluorescence intensity following 2-photon excitation^9^, was selected as FRET donor and the Cy3 derivative DiI was selected as FRET acceptor. To ensure efficient encapsulation inside LNDs with minimal dye leakage, DiI was coupled to tetraphenylborate (TPB)^10^ (**Figure 1B; Suppl. Figure S1 A**). The LNDs thus function as a FRET-based cargo delivery detection system (FedEcs). In contrast to previously reported FRET-LNDs operating in NIR region for whole mice imaging^7^, here we propose FRET-LNDs specially designed for 2-photon intravital imaging. This configuration enables targeted three-dimensional (3D) imaging of nanoparticles circulating within the brain vasculature at micrometer spatial resolution and sub-second temporal resolution. LNDs with diameter 30- and 80-nm, as measured by dynamic light scattering, containing either the donor, acceptor, or both dyes were formulated (**Suppl. Figure S1B**). At an excitation wavelength of 380 nm, LNDs of both sizes showed the fluorescent properties of the donor, *i*.*e*. fluorescence emission at 450 nm, as well as a second fluorescence peak at 580 nm, corresponding to the expected emission of the acceptor due to FRET (**Suppl. Figure S1 C**,**D**). The composition of the generated LNDs consisted of 98% lipids, with the remaining 2% accounted for by the donor and acceptor fluorophores. At this weight, FRET efficiency between the donor and acceptor reached 86% for 30- and 80-nm LNDs (**Suppl. Figure S1 E**), indicating a highly effective energy transfer between the fluorophores. Dilution of FedEcs-LNDs into dioxane, a solvent known to disrupt LND structure, led to the loss of FRET, as indicated by a fluorescence emission dominated by the donor dye alone (**Figure 1C**). These data demonstrate that our FRET system can reliably detect LNDs integrity and monitor cargo delivery.

**Figure 1.**
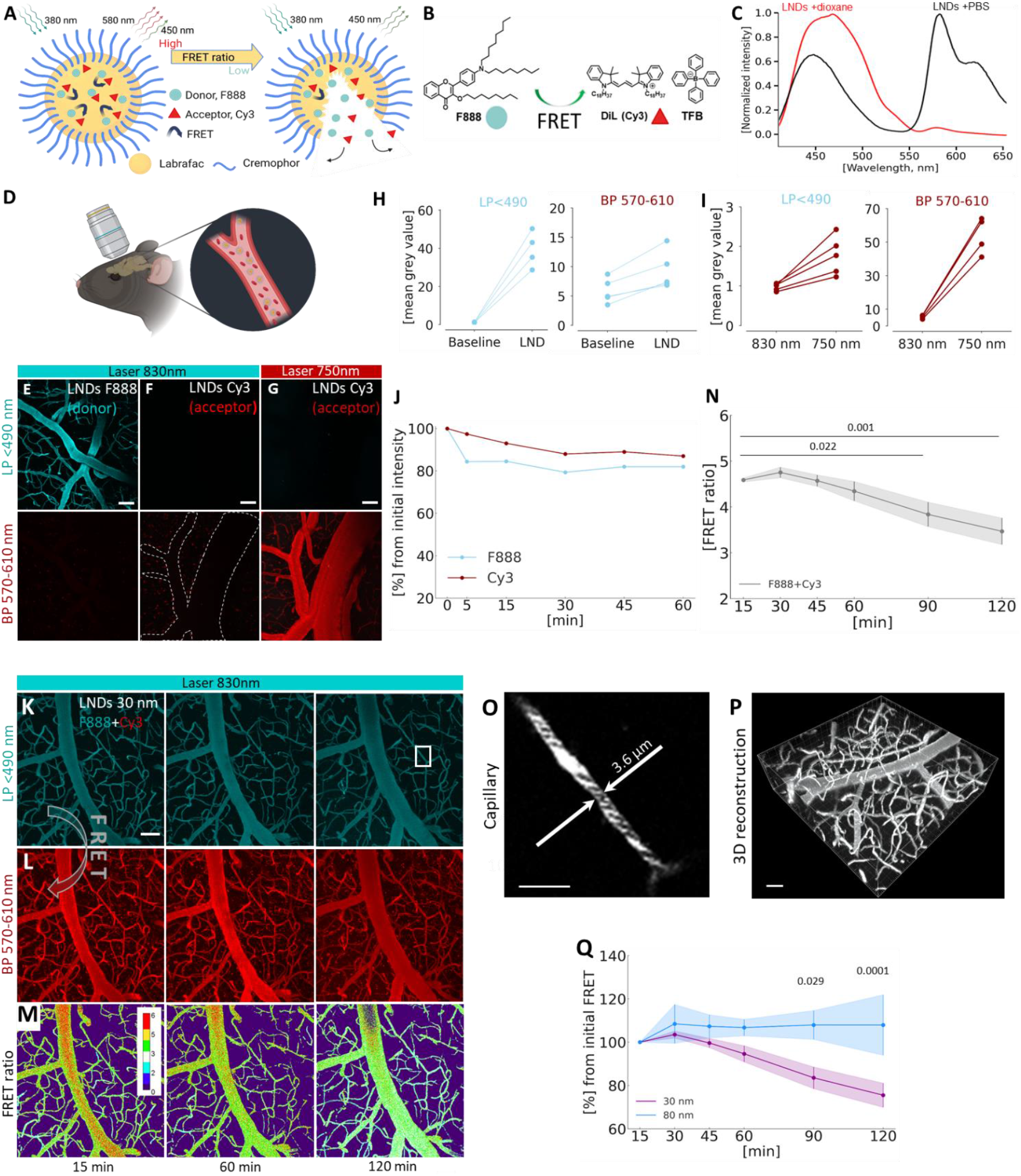
Design, characterization, and in vivo evaluation of lipid nanodroplets (LNDs) for real-time monitoring of drug release. **A**. Schematic representation of Förster-resonance energy transfer (FRET) nanocarrier that can shift its color upon decomposition, thus reducing the total FRET ratio. The LND consists of an oil core (yellow; Labrafac WL), surfactant (blue; Cremaphore, Kolliphor® ELP), and the fluorescent FRET pair dyes F888 (blue sphere) and Cy3 (red triangle) bound to a hydrophobic counter-ion. **B**. Chemical structure of F888 and Dil (Cy3) coupled with a counterion tetraphenylborate, responsible for causing FRET effect. **C**. *In vitro* fluorescence spectrum of FRET-LNDs containing F888 and DiI-TPB (2 wt% loading each) diluted in PBS (back) and dioxane (red). **D**. Schematic drawing of intravital imaging of mouse cortical vessels performed via cranial window in the skull. **E-G**. Representative *in vivo* 2-photon images of mouse cortical vessels after injection of 30-nm LNDs loaded solely by F888 (donor; excitation laser wavelength 830 nm; **E**) or Cy3 (acceptor; excitation laser wavelength 830 nm and 750 nm; **F** and **G**). Maximum intensity projection (MIP) of 150 µm Z-stacks; laser power 4.5-13%; donor channel – LP<490 nm; acceptor channel – BP 570-610 nm; scale bar - 50 µm. **E**. Bright fluorescent signal was observed in donor channel at 830 nm. **F**. Absence of a signal in both the donor and acceptor channel. Dashed line outlines invisible vessel, which become visible only at 750 nm. **G**. Bright fluorescent signal was observed in acceptor channel. **H**. Quantification of increased fluorescence intensity after injection of donor-loaded LNDs (n=5 regions of interest in the vessels) in the donor (LP<490) and acceptor (BP 570-610 nm) channel. **I**. Fluorescence intensity increase observed in acceptor-loaded LNDs after shifting the laser wavelength from 830 nm to 750 nm in both the donor and acceptor channel (n=5 regions of interest in the vessels). **J**. Elimination rates of LNDs loaded by F888 (donor) and Cy3 (acceptor) from the blood circulation of naïve mice during 1 hour (n=1). **K**,**L**. Representative real-time *in vivo* 2-photon MIP images of mouse cortical vessels captured during 15-120 min after injection of 30-nm FedEcs-LNDs loaded by FRET pair (F888+Cy3), excitation laser wavelength 830 nm. The acceptor became visible via BP 570-610 detector due to the FRET effect at an excitation laser wavelength of 830 nm. Scale bar - 50 µm. **M**. Color-code of FRET ratiometric changes in circulated FRET-LNDs. Scale bar - 50 µm. **N**. Quantification of time-depended LNDs disintegration in the mouse brain vasculature measured by changes in FRET ratio. Data are presented as mean ± standard deviation (SD). One-way ANOVA was used, n=3. **O**. High resolution image showing distribution of LNDs within capillaries and between circulated blood cells. Scale bar - 20 µm. **P**. 3D reconstruction of real-time imaging of circulated LNDs in the brain vasculature. Scale bar - 50 µm. **Q**. Comparative analysis of the degradation of circulated LNDs of two different sizes (30- and 80-nm) using FRET ratio. Data presented as mean ± SD. Two-way ANOVA was used, n=3. Created with BioRender.com.

### Using FedEcs to monitor the stability of circulating LNDs in healthy brain vessels *in vivo*

To investigate the critical question of whether FedEcs-LNDs are bright enough to be visualized *in vivo* and to validate their functionality, we injected FedEcs-LNDs systemically into mice and investigated the cerebral vasculature **(Figure 1D)** by *in vivo* 2-photon microscopy (2-PM). The FedEcs-LNDs exhibited strong fluorescence, allowing clear visualization in vivo. As expected, LNDs loaded with either the donor (F888) or acceptor (DiI-TPB) fluorophores emitted fluorescence only when excited at the specific wavelength for each fluorophore (830 nm for F888 and 750 nm for DiI-TPB; **Figure 1E-I**). Longitudinal imaging confirmed the stability of the circulated LNDs *in vivo* loaded with either fluorophore individually (**Figure 1J**). LNDs loaded with both FRET donors and acceptors (FedEcs-LNDs) emitted bright fluorescence at 830 nm in both channels (**Figure 1 K, L**), corroborating the *in vitro* data and confirming that FRET occurs in vivo. This finding suggests that FRET can be dynamically assessed in real time, allowing us to monitor particle integrity by quantifying the FRET ratio, expressed as the acceptor-to-donor fluorescence intensity ratio (**Figure 1M**). Real-time analysis of the FRET ratio of circulating 30-nm LNDs in healthy mice revealed a slow decrease in the FRET signal, which dropped from 4.6 to 3.5 of FRET ratio value over 120 min (**Figure 1N**). Notably, the spatial resolution of monitored FedEcs-LNDs was enough to observe motion of nanoparticles within the capillary and between blood cells and across the 3D volume of the brain (**Figure 1 O,P**). Interestingly, 80-nm LNDs had a substantially higher (p=0.0001) stability in the blood circulation, maintaining intact structures for two hours, while 30-nm lost ∼20% of integrity **(Figure 1Q)**. This enhanced stability is likely due to the higher oil-to-surfactant ratio in the larger particles. These findings demonstrate that our system can dynamically assess the real-time integrity of circulating nanoparticles using the FRET ratio. To our knowledge, this is the first report of a nanoparticle system able to produce a stable FRET signal *in vivo*, suitable to longitudinal high resolution intravital 2-PM.

### Employing FedEcs to define the distribution of LNDs after ischemic stroke *in vivo*

To investigate the behavior of LNDs in the brain under pathological conditions, we induced focal cerebral ischemia by occluding the middle cerebral artery of a mouse with an endovascular filament for one hour and systemically injected highly fluorescent LNDs 30 min after reperfusion (**Figure 2 A; Suppl. Figure S2A-B**). We focused on 30 nm LNDs, due to their prolonged blood circulation and optimal size, which enables them to cross the BBB via transcytosis, as shown in our recent study^11^. Brain tissue was harvested 60 minutes after LND injection and we detected a wide spread accumulations of LNDs within the ischemic territory (**Suppl. Figure S2C**). Staining of blood vessels (lectin) and red blood cells (Ter119) demonstrated that LNDs were located within cerebral capillary and colocalized with stalled erythrocytes. Interestingly, while conventional fluorescent tracer Cascade Blue (3000 Da), co-injected with LNDs, was diffusively distributed in the lesion area, the distribution of LNDs had a specific pattern: the intensity peak inside of microclots and a high intensity area in proximity of the vessel occlusions (**Figure 2B; Suppl. Figure S2C, i-iii**). This implicate that 30-nm LNDs may passively accumulate inside microthrombi and extravasate into the brain parenchyma.

**Figure 2.**
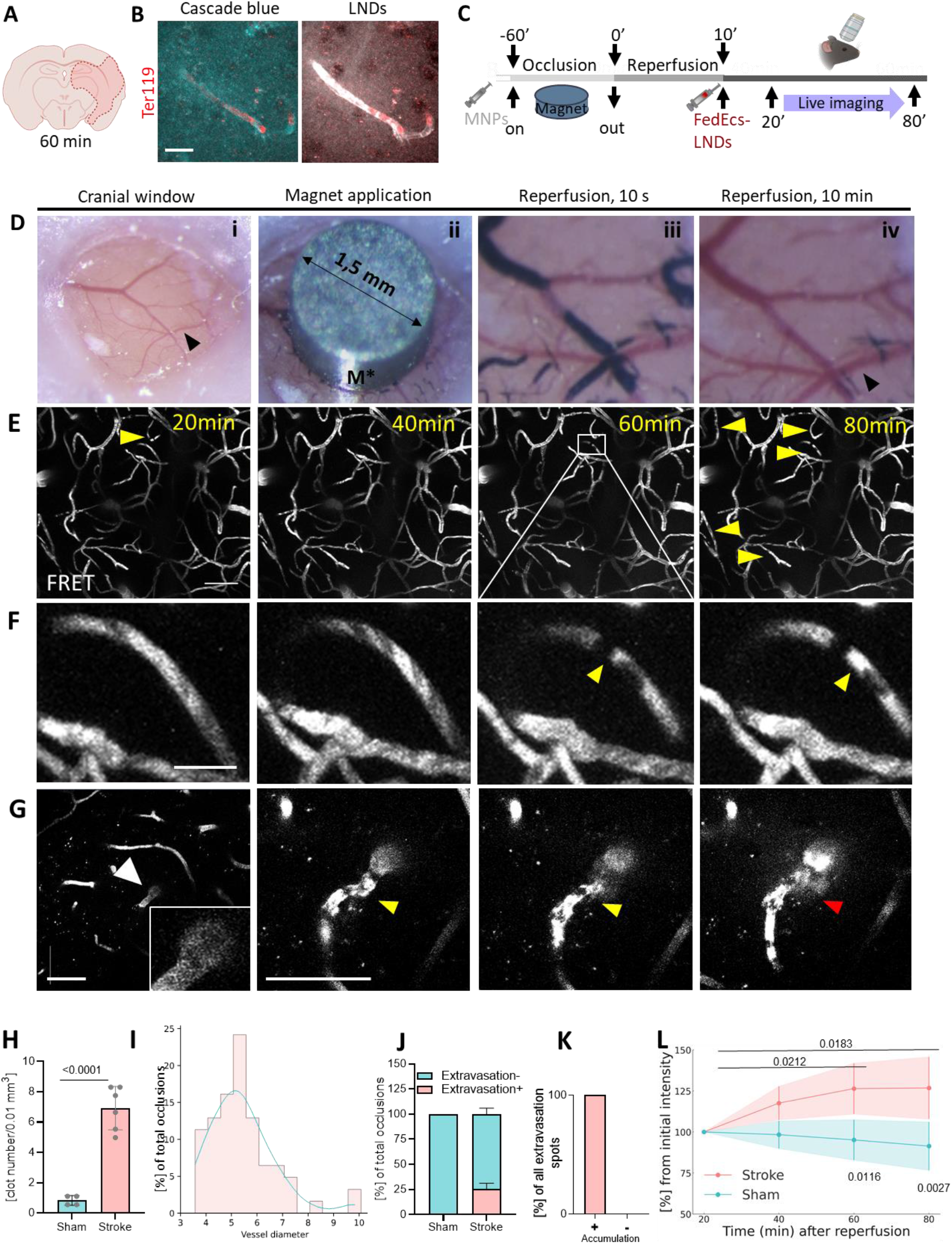
Spontaneous post-stroke microclot formation opens the blood-brain barrier for LNDs. **A**. A filament middle artery occlusion (fMCAo) during 60 min followed by injection of 30-nm LNDs loaded with rhodamine were performed as described at (**Suppl. Fig S2A-B). B**. Representative confocal image of the stroke-affected area in a mouse brain, showing extravasation in the microclot region. Stalling erythrocytes in the microclot area are stained with the Ter119 antibody (red), allowing comparison of the extravasation of Cascade Blue 3000 Da dextran *vs*. LNDs. Scale bar – 20 µm. **C**. Nano-stroke model experimental design. A mini magnet was placed for 60 min on the top of the glass-covered cranial window. Magnetic nanoparticles (MNPs) were injected systemically causing the occlusion of the vessels underneath the magnet. 20 minutes after removal of the magnet, the animal was placed under the 2-photon microscope (2-PM) for live imaging. **D**. Macrophotographs of different stages of the stroke model. i - glass-covered cranial window; black arrow – middle cerebral artery (MCA). ii - micro-magnet (M*; diameter 1.5 mm) placed on top of the cranial window located above the territory of the MCA. iii - MNPs accumulated in the vessel lumen of cerebral vessels underneath the magnet (black). iv - vessels re-perfused within 10 min after removal of the magnet; black arrow – MCA. **E**. Intravital 2-PM images of cortical vessels of the mouse with Nano-stroke model injected with FedEcs-LNDs. Yellow arrows – spontaneous vascular occlusions. Scale bar – 50 µm. **F**. Zoomed from **E**., a representative real-time images of spontaneous microvascular clot formation with subsequently accumulated LNDs (yellow arrow). Scale bar – 10 µm. **G**. Representative longitudinal real-time images of spontaneous microvascular clot formation followed by extravasation of LNDs into the brain parenchyma (yellow arrow). Scale bar - 50 µm. **H**. Analysis of numbers of microvascular clots at 80 min post-reperfusion compared to sham animals. Data are presented as mean ± standard deviation (SD). Unpaired t-test. Stroke: N=79 clots (6 mice); Sham – 4 mice. **I**. Size-distribution of the vessels with microclots. Data are presented as proportion of vessel diameter where microclots were detected. N=62 clots (6 mice). **J**. Analysis of association between microclot (accumulation) and blood-brain barrier dysfunction (extravasation). N=20. **K**. Proportion of microvascular clots associated with extravasation of the LNDs. Data represented as mean ± SD. N=4 (sham) and n=6 (nano-stroke). **L**. Longitudinal analysis of the accumulation of FedEcs LNDs in the ischemic brain area. Data represented as mean ± SD. Two-way ANOVA was used, n=4 (sham) and n=6 (nano-stroke). Created with BioRender.com.

To further investigate this phenomenon, we then aimed to analyze LNDs biodistribution real-time and with the highest possible resolution to decipher the route/mechanism by which LNDs extravasated and whether the accumulation of LNDs in ischemic brain may have affected their ability to carry and deliver cargo. For this purpose, we developed a novel stroke model based on a technique previously used for inducing focal cerebral ischemia in juvenile mice^12^. This model employs iron oxide magnetic nanoparticles (MNP) to transiently occlude cortical blood vessels. Following occlusion, the animal is placed under 2-PM for *in vivo* imaging of the affected cortical area **(Figure 2C)**. In detail, animals received a systemic injection of MNP and subsequently a strong mini-magnet was placed on top of implanted cranial window to occlude vessels at MCA area (**Figure 2D, i**). The magnetic field trapped the 180 nm pegylated MNPs within the cerebral vasculature (**Figure 2D, ii**) thereby causing cerebral ischemia (Nano-Stroke). After one hour, removing the magnet allowed the MNPs to wash away, mimicking the recanalization after stroke and allowing the tissue to be reperfused (**Figure 2D, iii-iv**). Using EM, we were able to resolve single iron oxide particles and demonstrate that, during occlusion, MNPs caused localized astrocytic swelling **(Suppl. Figure S3A)** without damaging the vessel intima **(Suppl. Figure S3B)**, indicating ischemic tissue damage. This model occluded cortical blood vessels up to a depth of 500 µm (**Suppl. Figure S3C**) and allowed immediate live 2-PM of reperfused cerebral vessels (**Suppl. Figure S3D**). Histological evaluation of the lesion area demonstrated inflammatory reaction and neuronal loss (**Suppl. Figure S3 E-I**). Using the Nano-Stroke model and FedEcs-LNDs we investigated the passive biodistribution of systemically applied LNDs in the brain after experimental stroke by *in vivo* 2-PM focusing first on LNDs with preserved integrity according to their FRET (acceptor) signal (**Figure 2E**). Post-reperfusion, we observed LND accumulations within non-flowing segments of cerebral vessels increasing during imaging period (**Figure 2E, yellow arrows**). This accumulations with non-moved black cell shadows we consider to define as microvascular occlusions or microthrombi. Important to note, that some LND accumulations were newly formed or spontaneous **(Figure 2F, yellow arrow)**, while some also were accompanied by subsequent extravasation of LNDs into the brain parenchyma (**Figure 2G, 80 min, red arrow**). Quantification showed that the number of LND accumulations 80 min after ischemic stroke was 8-fold higher than in sham operated mice **(Figure 2H;** p<0.0001**; Suppl. Figure Fig. S4)** and that 90% of all observed occlusions occurred in vessels less than 6 µm, *i*.*e*. brain capillaries (**Figure 2I**), further implying that LNDs were trapped in microvascular-occlusions. Interestingly, 25% of all microthrombi observed after stroke were associated with extravasation of LNDs **(Figure 2J)**, while no extravasations were detected in sham operated animals. Most importantly, extravasation of LNDs were only observed at sites where micro-occlusions formed **(Figure 2K)**, suggesting that micro-occlusions compromised the permeability of the adjacent capillary wall and were critically involved in the opening of the BBB. Longitudinal *in vivo* imaging of LNDs allowed us to detect that particles gradually accumulated within the ischemic brain (**Figure 2L**). Altogether, following ischemic stroke, 30-nm LNDs passively accumulated within the vessel lumen around the microclot over 80 minutes, and also extravasated from there into the ischemic brain parenchyma. Our data provide the first visualization of LNDs entering the ischemic brain via microthrombi, implicating new avenue to deliver drugs into ischemic tissue.

To better understand local distribution of LNDs inside occluded vessel as well as morphology and ultrastructure of vascular occlusions and extravasation sites, we developed novel correlative light-electron microscopy (CLEM) method. Since MNPs are both visible at light microscope and electron-dense, they can be used as fiducial markers for CLEM. Capitalizing on this feature, we tracked fluorescent emission of LNDs accumulations and extravasations in real-time using 2-PM *in vivo*. We then correlated these dynamic observations with ultrastructural data by relocating the fluorescent signal to EM and performing 3D array tomography via automated tape-collecting ultramicrotomy (ATUM) to generate ultrastructural 3D images of the chosen clot **(Figure 3A; Suppl. Figure S5; Methods**). On 2-PM time-serial Z-stacks of a mouse brain after 60 min stroke, we located two representative microvascular occlusions marked by LND-FedEcs accumulation appeared at different time points—20 minutes and 80 minutes after reperfusion **(Figure 3B, yellow and magenta arrows)**. Then occlusions were relocated using CLEM and successfully re-identified at EM **(Figure 3C; Movie S01)**. After full reconstruction of the vessel segments in 3D (**Figure 3D**; **Movie S02**), we could pinpoint the cellular composition and the microstructure of the occluded vessels at the LNDs accumulation point (**Figure 3E-F**). Microglia and constricted pericytes surrounded the occluded vessels, while erythrocytes, platelets and LNDs co-localized within the occlusions (**Figure 3E-F**). When zooming on the ultrastructure of the microvascular occlusions, we found fibrin nets (orange arrow) in between red blood cells, further suggesting that the observed occlusions were clearly identified as microthrombi (**Figure 3E-F, i**). Interestingly, microclots which formed early after reperfusion according to *in vivo* imaging contained erythrocytes with affected morphology, while those formed later had erythrocytes with the normal biconcave shape (**Figure 3E-F, ii, red**). Collectively, these findings show that stroke induces the formation of microthrombi within the cerebral microcirculation and that LNDs trapped within these microclots independent of their time formation.

**Figure 3.**
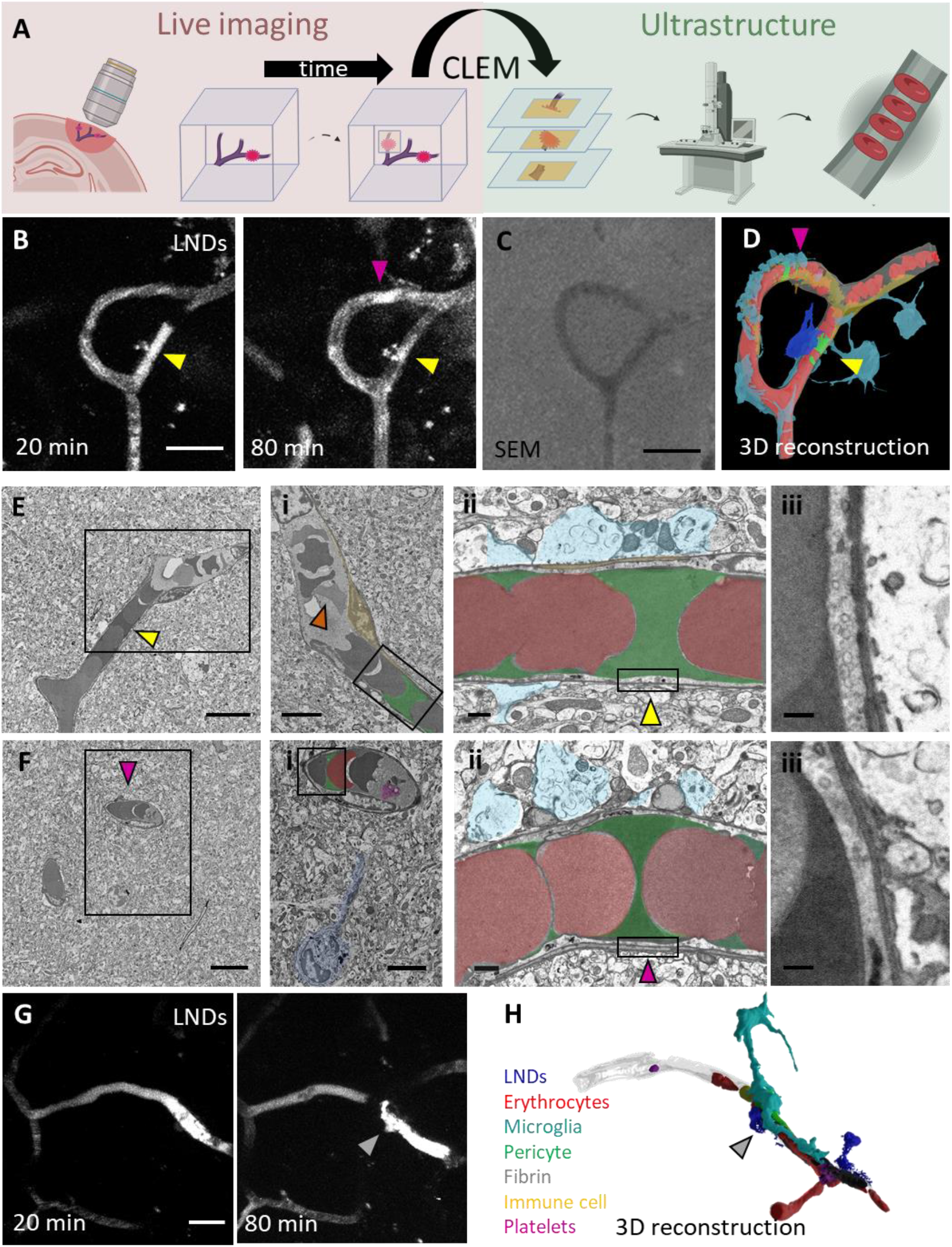
Correlative light-electron microscopy revealed ultrastructural changes in sites of accumulation and extravasation of FedEcs-LNDs. **A**. Experiment design as described in **Suppl. Fig. S3A** with longitudinal intravital imaging during 60 minutes followed by fixation, embedding and serial sectioning for correlated volume electron microscopy (EM) by automated serial sections to tape (ATUM; **Suppl. Fig. S5**). **B**. Representative 2-PM images of post-stroke mouse brain capillaries containing “old” (20 min post-reperfusion; yellow arrow) and “fresh” (80 min; magenta arrow) spontaneous microvascular clots. Scale bar 20 µm. **C**. Representative example of a microvascular clot (20 min) transitioning towards LNDs extravasation (80 min; grey arrow) into the brain parenchyma (“leaky” microclot). Scale bar 20 µm. **D**. Summed slices of a **s**canning electron microscopy (SEM) stack of the relocated region from the 2-PM experiment **(Suppl. Fig.S5)**. Scale bar 20 µm. **E**. Rendered three-dimensional reconstruction of the volume SEM data thereof. Red (erythrocytes); yellow (pericytes); green (LNDs); sky blue (astrocytes); dark blue (microglia); magenta (platelet). **F**. Rendered three-dimensional reconstruction of volume electron microscopy of the relocated region from **Fig. 3C**. Red – erythrocytes; dark blue – LNDs; cyan – microglia; grey – fibrin; green – pericyte; magenta – platelets; yellow – leukocytes; transparent grey – BEC. See in Movie S03. **G.-H**. Single segmented high-resolution SEM images of the “old” (G; yellow arrow) and “fresh” (H; magenta arrow) microclots from 3B. Scale bar 10 µm. **G**,**i**. Zoomed area from G. containing an “old” microclot with accumulated LNDs (green) and a pericyte constricting the capillary (yellow). Orange arrow – dysmorphic erythrocytes trapped into fibrine. **G**,**ii**. Zoomed area from E,i containing an occluded capillary with accumulated LNDs (green) between erythrocytes (red). Sky blue – astrocytic end feet. Yellow arrow – relocated capillary area with accumulated LNDs showing a higher than in normal capillaries fluorescent intensity in 2-PM. Scale bar – 1 µm. **G**,**iii**. Zoomed brain endothelium cell (BEC) from J,ii. containing a large number of vesicular-like structures. Scale bar – 200 nm. **H**,**i**. Zoomed area from H. containing a “fresh” microclot with accumulated LNDs (green) between erythrocytes (red) and microglia cell (dark blue) with its processes directed towards the platelet cells (magenta) trapped into fibrine. **H**,**ii**. Zoomed area from H,i containing an occluded capillary with accumulated LNDs (green) between erythrocytes (red). Sky blue – astrocyte end feet. Magenta arrow – relocated capillary area with accumulated LNDs having higher fluorescent intensity in 2-PM. Scale bar – 1 µm. **F**,**iii**. Zoomed BEC from F,ii. with low level of caveolae-like structures. Scale bar – 200 nm. **H**,**iii**. Zoomed BEC from H,ii. with high level of caveolae-like structures. Scale bar – 200 nm. **I**. Zoomed F. demonstrating that extravasation occurs from the place where microthrombi were formed. FedEcs LNDs – dark blue; Erythrocytes – red; Microglia – cyan; Pericyte – green; Fibrin – grey; Immune cell – yellow; Platelets – magenta. Grey arrow – place of extravasation relocated from **Fig. 3C**. Created with BioRender.com.

Using CLEM, we precisely investigated the microstructure of brain endothelial cells (BECs) in the proximity of microclots with accumulated LNDs (**Figure 3E-F, ii-iii**). The BECs in the vicinity of older clots had a much higher number of caveolae (23 caveolae/µm; **Figure 3E, iii**) than BECs next to fresh microclot (2.8 caveolae/µm; **Figure 3F, iii**), implicating that a persisting contact between BECs and clots may induce the formation of caveolae, the prerequisite for the subsequent compromising of BBB and, as a result, extravasation of LNDs into the brain parenchyma. Additionally, a representative microclot that became “leaky” 60 minutes after initial occlusion, detected by LND accumulation **(Figure 3G, grey arrow)**, provides clear evidence that the persistence of microclots compromises the BBB in their proximity. To confirm it, we performed the CLEM for the representative “leaky” clot with accumulated LNDs and extravasation next to it. A 3D reconstruction of the ultrastructure revealed that the extravasation site **(Figure 3H, grey arrow)** was fully co-localized with microthrombi composed of erythrocytes, platelets, and immune cells **(Movie S03)**. Outside the vessel, the microclot was surrounded by pericytes and microglial processes, resembling the inner and outer composition of two other above-mentioned reconstructed clots.

Thus, our electron microscopy data demonstrate that stroke-induced persistent microclots within the cerebral microcirculation progressively create an environment conducive to increased endothelial transcytosis via caveolae, which may be associated with BBB disruption and the subsequent entry of LNDs into the brain parenchyma.

To investigate this process on the functional level *in vivo*, mice received a systemic injection of LNDs first and were only then subjected to occlusion of the vessels via magnet-MNP interaction for 60 min. The cerebral microcirculation was imaged by *in vivo* 2-PM before and after occlusion **(Suppl. Figure S6)**. Images obtained immediately after recanalization showed multiple spots of LND extravasation at the occluded area **(Suppl. Figure S6, yellow arrows)**, supporting our histological and EM findings that micro-occlusion of the vessels (even caused by iron oxide nanoparticles) may serve as entry point for LNDs into the brain parenchyma. Thus, by occluding vessels with MNPs, we were able to locally open the BBB.

### Monitoring cargo delivery of LNDs in the brain using FedEcs

The entry of LNDs into the brain parenchyma is, however, not equivalent with cargo delivery. Therefore, we used the newly developed FedEcs functionality of our highly fluorescent LNDs to monitor whether LNDs decompose after being trapped in microclots and transmigrate into the brain parenchyma. To assess this, we performed a nano-stoke **(Figure 4A)** and measured the FRET ratio in flowing versus non-flowing segments of the microvasculature after cerebral ischemia over time by quantitative *in vivo* FRET imaging **(Figure 4B)**. While FRET ratio of LNDs circulating in plasma was stable during 80 min of observation, the FRET ratio of LNDs trapped within microclots was significantly lower and gradually decreased **(Figure 4C)**. The loss of FRET is caused by a loss of interaction between the two fluorophores undergoing FRET and is therefore a quantitative measure of particle decomposition. Hence, our data indicate that LNDs are stable in plasma, but start decomposing once being trapped in microclots. This observation is further confirmed by the analysis of newly formed individual microclots over time **(Figure 4D)**. Once trapped in a formed microclot, the FRET ratio of LNDs starts declining and reaches about 50% of its baseline value after 40 min. Interestingly, a very similar trend to decay of FRET ratios can also be observed when focusing the analysis on LNDs locally extravasated into the brain parenchyma **(Figure 4E)**. The presence of a FRET signal in the parenchyma indicates that intact LNDs entered the brain via microthrombi. Additionally, these LNDs show a constant decay of their FRET ratio suggesting that LNDs release their cargo, in our case the fluorophores undergoing FRET, after entering the brain parenchyma.

**Figure 4.**
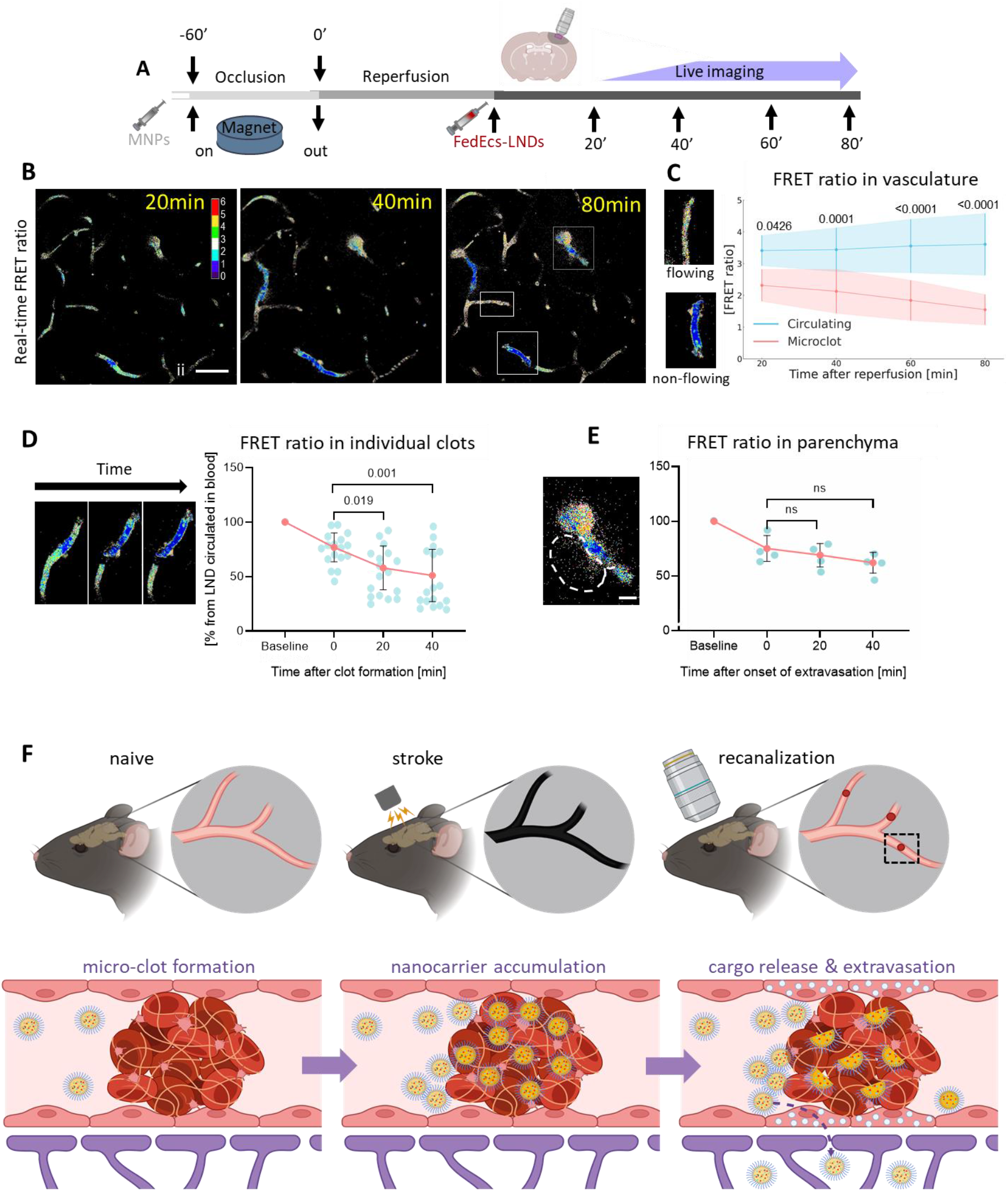
FRET ratio reveals higher decomposition of FedEcs-LNDs in microclots, but not in brain parenchyma as well as delivery of cargo into the brain parenchyma. **A**. Experiment design. **B**. Representative longitudinal real-time images of visualization of FRET ratio related to decomposition of FedEcs-LNDs in vasculature of ischemic brain area. Scale bar - 50 µm. **C**. Longitudinal analysis FRET ratio related to decomposition of FedEcs-LNDs in vasculature of ischemic brain area during 80 minutes after recanalization. Data are presented as mean ± SD. Two-way ANOVA was used, circulated n=16-17 ROIs; microclots n=6-13; 5 mice. **D**. Time-dependent analysis of relative FRET changes of LNDs located inside individual clot. **E**. Time-dependent analysis of relative FRET changes of extravasated into brain parenchyma LNDs. Data are presented as single points and mean ± SD. One-way ANOVA was used, n= 17 clots and n=4 extravasations, respectively. Scale bar – 10 µm. Yellow arrow – clot; dashed area – extravasated FedEcs-LNDs emitting FRET. Created with BioRender.com.

In summary, using highly fluorescent LNDs and FedEcs, a novel *in vivo* real-time cargo release detection system, we unraveled the passive biodistribution of LNDs after ischemic stroke *in vivo* and at ultrastructural level **(Figure 4F)**. Our data demonstrated that after transient ischemic stroke caused by MNPs-magnet interaction the newly formed microclots are developed in the stroke area. FedEcs lipid nanodroplets accumulated inside these microthrombi releasing their cargo there, crossed the BBB and entered the brain parenchyma. Thus, FedEcs represents a novel and valuable tool to investigate the therapeutic potential of nanoparticle-based drug delivery systems and to directly visualize cargo delivery *in vivo* with a high spatial and temporal resolution.

## Discussion

Targeted drug delivery using nanotechnology has two conflicting aims: nanoparticles must reliably enclose their cargo and transport it to the target tissue, but release it there effectively^13^. Both aims are hard to achieve and even more challenging to investigate experimentally due to insufficient technologies for particle tracking and cargo release in living organisms^2^. In the current study we developed highly fluorescent LNDs with a built-in cargo release function based on FRET and used these nanoparticles (NPs) to investigate the biodistribution of LNDs and cargo release in a novel, magnetic NPs-based model of ischemic stroke in mice. One of the key observations in our study was the passive accumulation of 30-nm LNDs within newly formed microclots following transient ischemic stroke. This size of LNDs was specifically chosen due to their demonstrated ability to cross the BBB via transcytosis, as shown in our previous work^11^. Importantly, our data showed that LNDs not only accumulated within the ischemic territory but also crossed the compromised BBB, entering the brain parenchyma. This is significant as it suggests that microclots may serve as an entry point for nanoparticles, such as LNDs, potentially allowing the delivery of therapeutic agents directly to affected brain tissues. We confirmed this using MNPs-based microclots to open the BBB. Additionally, our data explain the cellular and subcellular mechanisms responsible for the biodistribution of lipid nanoparticles in ischemic brain published earlier^14-16^, define the exact site of passive particle retention, and suggest LNDs as potential drug carriers for the delivery of therapeutics to the brain under conditions where microthrombi are present, *i*.*e*. ischemic stroke or traumatic brain injury^11^.

To investigate the dynamic and ultrastructural insights of biodistribution of systemically applied NPs in the brain after transient stroke, we had to develop and improve several novel technologies. After observing retention of LNDs inside microclots in the ischemic brain by confocal microscopy, we used *in vivo* deep brain 2-photon imaging to understand the dynamics and the mechanism of this process. For this purpose, we developed a new MNPs-based stroke model, nano-stroke, which allowed us to image exactly the same cerebral vessels before and after reperfusion. The nano-stroke model is based on the recently published SIMPLE stroke model performed in pups^12^. We adapted the SIMPLE model to adult animals by implanting a chronic cranial window, thus allowing us to visualize cerebral vessels in the ischemic brain longitudinally and with high resolution. Combining the nano-stroke model with highly fluorescent, super-bright FedEcs-LNDs and intravital 2-PM, we were able to demonstrate real-time that following reperfusion systemically injected LNDs accumulate inside microthrombi and extravasate there, a so far missed phenomenon due to the use of low-fluorescence NPs. Further, the novel CLEM method we developed allowed us to match real-time 2-PM observations with ultrastructural data at a high spatial resolution. This methodology enabled us to visualize LNDs accumulation within microthrombi and correlate it with 3D structural features such as fibrin nets and erythrocyte morphology. Moreover, this technique enabled us to combine a dynamic component (timing of clot appearance) with ultrastructural information of BEC, thereby demonstrating that microclots compromise the function of the BBB by an increase formation of caveolae.

Accumulation of LNDs inside microclots after cerebral ischemia may have a large medical potential. The standard treatment for ischemic stroke is recanalizing large cerebral arteries either by dissolving or removing the occluding thrombus by systemic administration of recombinant tissue plasminogen activator or by mechanical thrombectomy, respectively^17^. However, in many cases recanalization of large cerebral vessels does not result in reperfusion of the cerebral microcirculation and restoration of cerebral blood flow and may thus allow additional tissue damage to occur^18^. In previous attempts to dissolve microthrombi after stroke researchers attached the main thrombus-attaching protein (Cys-Arg-Glu-Lys-Ala) CREKA on the surface of nanocarriers^19-21^, however, due to the lack of appropriate imaging techniques it remained elusive whether these particles delivered their cargo into microclots. In the current study, we did not use any active targeting strategy, still our LNDs ended up in large numbers within microclots. This passive biodistribution of LNDs in post-stroke microclots is of fundamental medical importance and may pave the way for future approaches to target microthrombi with respective active compounds.

It still remains unclear how such high numbers of NPs get trapped in microthrombi. One possible explanation could be that NPs get trapped in forming microthrombi, *i*.*e*. while the clot forms, however, the large number of NPs located between all components of the clot does not seem to support such a scenario. The more likely explanation is that it takes some time until clots form and fully occlude a cerebral microvessel, as suggested by our previous findings and by others^22-24^. During this period, which may last for minutes or even hours, vessels are not patent for cells but for plasma and since NPs travel with plasma, they enter the clot and get trapped in between accumulated fibrin fibers and stalled red and white blood cells. This “filtering” mechanism fully explains the large number of LNDs we found in microclots and the specificity with which LNDs accumulate in these structures. This concept is well in line with previous studies which showed neuroprotective effects with nano-scale compounds after ischemic stroke, but did not elaborate on the mechanisms of particle distribution and cargo delivery^25, 26^. The accumulation and subsequent extravasation of nanocarriers at the microclot site is particularly advantageous, as it halts their motion in the bloodstream, increasing the likelihood of interaction with BEC caveolae and enhancing the concentration of nanoparticles that cross the BBB, thereby maximizing their therapeutic potential in the brain parenchyma.

In the current study we chose LNDs as nanocarriers because they are regarded to be non-toxic and biodegradable and they have a relatively long half-life of up to 8h in mammalian blood^7^. To monitor cargo release, we equipped LNDs with a FRET-based cargo release detection system, FedEcs. FRET systems have been widely used in nanoscience for more than a decade^27^, however, the novelty of our current approach was to use a FRET pair compatible specifically with 2-photon excitation for high resolution intravital imaging of cargo release. This approach resulted in LNDs with such a high FRET signal that LNDs could be visualized by 2-PM at subcellular level in mouse blood and brain *in vivo*. In comparison to recent, more traditional approaches to image drug delivery to the brain using fluorescent NPs^14, 28, 29^, FedEcs has the large advantage of being able to visualize the actual moment when LNDs decompose and release their cargo, even in tiny amounts of crossed BBB LNDs. Thereby, FedEcs can clearly distinguish between artefactual cargo delivery to the brain—where dyes leak from circulating nanocarriers and cross the BBB—and factual cargo delivery, where the carrier itself crosses the BBB and releases its cargo thereafter. Thus, FedEcs will allow researchers to monitor the release of cargo from individual NPs with high spatial and temporal resolution in all organs accessible to *in vivo* microscopy avoiding all ambiguities caused by previous methods.

## Conclusion

In conclusion, we successfully developed highly fluorescent lipid nanodroplets carrying a FRET-based cargo delivery system (FedEcs). FedEcs-LNDs are bright enough to be visualized individually in blood and brain of living mice by 2-photon microscopy. Using in vivo brain imaging we were able to quantify the stability of LNDs in blood and demonstrate the preferential accumulation of LNDs in microvascular clots, extravasation of intact LNDs across the blood-brain barrier, and cargo delivery to the brain parenchyma following cerebral ischemia. Therefore, FedEcs is a reliable and potent dynamic indicator of cargo delivery and release and provide a future testing ground for the efficacy and delivery of a wide-range of targeted nanoscale therapy in future.

## Acknowledgements

The authors thank Georg Kislinger and Hanyi Jiang for the segmentation and rendering of the 3D EM model and Platin Ramadani for the rendering. This work was supported by the European Union Horizon 2020 research and innovation program under the Marie Skłodowska-Curie grant agreement No 794094, the European Research Council ERC Consolidator grant BrightSens 648528, the Agence National de Recherche ANR SenEmul ANR-22-CE18-0034, and funded by the Deutsche Forschungsgemeinschaft (DFG, German Research Foundation) – projects number: 457586042, FOR2879; A03 – Mi 694/9-1 – 428663564; DFG under Germany’s Excellence Strategy within the framework of the Munich Cluster for Systems Neurology (EXC 2145 SyNergy—ID 390857198) and the TRR 274/1 2020 – 408885537 (project Z01).

## Authors contribution

I.K. initiated and co-fund the study, led scientific aspects of the project, designed, performed and analyzed most of experiments, wrote the first draft of the manuscript. N.A. formulated and optimized FRET-LNDs for 2-photon imaging. M.S. coordinated and performed CLEM experiment, helped write the manuscript. A.W. performed and designed *ex vivo* part of Nano-stroke experiment and its analyses, helped write the manuscript. V.B.T. formulated FRET-LNDs and performed their analyses. U.M. performed fMCAo stroke experiment. J.S. performed Nano-stroke and fMCAo experiments and analysis. T.M. led EM part of the project, coordinated CLEM experiments and analysis. SF designed, coordinated, performed Nano-stroke and CLEM experiments. A.K. led FRET-LNDs aspect of the project, provided infrastructure, supervised and cowrote final draft of the manuscript. N.P. led main animal experimental aspects of the project, co-fund the study, provided infrastructure, supervised, cowrote the final manuscript. All authors critically reviewed, contributed to and approved the final manuscript.

## Material and Methods

### 1. Formulation of LNDs

Dye loaded nanoemulsions were obtained by spontaneous nanoemulsification. Briefly, the dyes R18-TPB (prepared as described before)^30^ was dissolved in LabrafacWL® at 2% by weight. In case of FRET-LNDs, F888 (FRET donor, prepared as described before^7, 9^) and DiI-TPB (FRET acceptor, prepared as described before^10^) Then, 60 mg of the oil with the dye were mixed with 40 mg Cremophor ELP® (also called Kolliphor ELP®) and homogenized under magnetic stirring at 40 °C for 10 min. The LNDs were obtained by the addition of ultrapure (Milli-Q) water (230 mg) under rapid magnetic stirring. Size distributions were determined by dynamic light scattering using a Malvern Zetasizer ZSP.

### 2. Animals

Male 6-8 weeks old, 18-22 g, C57Bl6/J mice from Charles River Laboratories (Sulzfeld, Bavaria, Germany) were used. Mice were group-housed under pathogen-free conditions and bred in the animal housing facility of the Institute of Stroke and Dementia Research (Muenchen, Germany), with food and water provided ad libitum (21 ± 1°C, at 12/12 hour light/dark cycle). Animal husbandry, health screens, and hygiene management checks were performed in accordance with Federation of European Laboratory Animal Science Associations (FELASA) guidelines and recommendations^31^. All experiments were carried out in compliance with the National Guidelines for Animal Protection of Germany with the approval of the regional Animal care committee of the Government of Upper Bavaria, and were overseen by a veterinarian. The data were collected in accordance with the ARRIVE guidelines^32^.

### 3. Stroke fMCAo

Ischemic stroke animal model via filament middle-cerebral artery occlusion (fMCAo) was performed as previously described^33^. Briefly, buprenorphine (0.1 mg/kg) injected animals, 30 min later were anesthetized with 1.8–2% isoflurane in 50% O2 in air by face mask under the control of rectal temperature (37°C±0.1°C) with a feedback-controlled heating pad. Regional cerebral blood flow (rCBF) over the territory of the middle cerebral artery (MCA) was monitored via glued onto parietal skull probe using laser Doppler fluxmetry (Perimed, Stockholm, Sweden). A silicone-coated monofilament (Doccol Corporation, Sharon, MA, USA) was introduced into the left common carotid artery and advanced toward the Circle of Willis until a drop of rCBF more than 80% of baseline indicated occlusion of the MCA. Thereafter, animals were allowed to wake up. After 60 min, animals were re-anesthetized and the filament was removed to allow reperfusion. Immediately, femoral artery catheter was inserted and 30 min post-reperfusion LNDs or MNPs were injected in volumes 4 or 6 μL/g per animal, respectively.

### 4. Ex vivo: confocal (fMCAo, nano-stroke)

Animals were injected via femoral catheter by 50 µl of DyLight 649 Labeled Lycopersicon Esculentum (Tomato) Lectin (Vector Laboratories, Burlingame, CA, US) and 5 min later were transcardially perfused with 4% PFA in deep anesthesia. Free floating coronal 80 µm sections were prepared as previously described^34^. The sections were blocked and simultaneously stained with the primary antibody in buffer (1% bovine serum albumin, 0.1% gelatin from cold water fish skin, 0.5% Triton X-100 in 0.01 M PBS, pH 7.2–7.4) for 72h at 4°C. The following primary antibodies were used: iba-1 (rabbit, Wako, #019-19741, 1:200), NeuN (guinea pig, Synaptic Systems, #266 004, 1:100), GFAP-Cy3 (mouse, Sigma Aldrich, #2905, 1:200). After incubation sections were washed in PBS and incubated with the following secondary antibodies: anti-rabbit coupled to Alexa-fluor 488 (goat anti-rabbit, Thermo Fisher Scientific, 1:100), anti-rabbit coupled to Alexa-fluor 594 (goat anti-rabbit, Thermo Fisher Scientific, #A-11012, 1:200), anti-guinea pig coupled to Alexa-fluor 647 (goat anti-guinea pig, Thermo Fisher Scientific, #A-21450, 1:200) in secondary antibody buffer (0.05% Tween 20 in 0.01 M PBS, pH 7.2–7.4). Nuclei were stained with 4′,6-Diamidin-2-phenylindol (DAPI, Invitrogen, #D1306) 1:10,000 or DRAQ5 (ThermoFisher, #65-0880-92) 1:1000 in 0.01 M PBS. Imaging was performed using confocal microscopy (ZEISS LSM 900, Carl Zeiss Microscopy GmbH, Jena Germany). For NeuN and iba-1 quantification, 40x magnification was used (objective: EC Plan-Neofluar 40x/1.30 Oil DIC M27) with an image matrix of 512×512 pixel, a pixel scaling of 0.4 × 0.4 μm and a depth of 8 bit. Three different ROIs in the lesion from superficial to deeper levels were chosen and collected in 15 μm z-stacks with a slice distance of 1 μm. Homotypical areas were imaged on the contralateral hemisphere and used as controls. MAP2 intensity was then measured and normalized to contralateral.

4.1 Analysis of microglia coverage. Microglia coverage was manually assessed in a maximum intensity projection of iba-1 stained sections. ROIs were chosen in the center of the lesion at 250 µm depth from the cortex surface at -1mm from bregma. Homotypical ROIs were chosen on the contralesional hemisphere. 3 ROIs per section per hemisphere were chosen. For microglia coverage, manual cell counting was used. To assess microglia morphology, Sholl and fractal analysis were performed to indicate ramification, cell range and circularity using a modified protocol from Young and Morrison^35^. Z-stack images were converted to a maximum intensity projection and microglia cells were individually cut out using the polygon selection tool in ImageJ. Only cells which were entirely contained within the z-stack were selected. Images were then thresholded and binarized as well as resized to 600 × 600 pixels, keeping the original scale. Any speckles and debris around the cell were removed using the paintbrush tool. Sholl analysis was performed using ImageJ ^36^. Centered on the soma, concentric circles with an increasing radius of 1 μm were drawn, the number of intersections measured at each radius.

After converting binary images to outlines, fractal analysis was performed using the FracLac plugin for ImageJ ^37^. As described previously ^35^, the total number of pixels present in the cell image of either the filled or outlined binary image were calculated and later transformed to μm2 (pixel area = 0,208 μm2). Cell circularity was calculated as

Circularity = 4*π*Area/Perimeter^2^.

#### Statistical analysis

All data is given as mean ± standard deviation (SD) if not indicated otherwise. For comparison between groups, Student t-test was used for normally distributed data and Mann-Whitney Rank Sum test for non-normally distributed data according to the result of Kolmogorov-Smirnov normality test. Measurements over time were tested between groups using One-way or Two-way ANOVA with Repeated Measurements, followed by Turkey’s multiple comparisons test for normally and Sidak’s multiple comparisons test for non-normally distributed data as post hoc test. Calculations were performed with Sigmaplot version 14.0 (Systat Software GmbH, Erkrath, Germany) and GraphPad Prism version 8.4.3 (GraphPad Software, San Diego, California USA).

### 5. Cranial window preparation

Before use, surgical tools were sterilized in a glass-bead sterilizer (FST, Heidelberg, Germany). Mice were anesthetized by an intra-peritoneal injection of medetomidine (0.5 mg/kg), fentanyl (0.05 mg/kg), and midazolam (5 mg/kg) (MMF) (140/10 mg/kg body weight, WDT, Bayer, Leverkusen, Germany). Subsequently, mice were placed and head-fixed in a stereotactic frame (David Kopf Instruments, Tujunga, CA, USA). Throughout the experiment, body temperature was monitored and maintained by a rectal probe attached to a feedback-controlled heating pad (Harvard Apparatus, Holliston, MA, USA). Eyes were protected from drying by applying eye ointment (Bepanthen, Bayer, Leverkusen, Germany). The scalp was washed with swabs soaked with 70 % ethanol. A flap of skin covering the cranium was excised using small scissors. The periosteum was scraped away with a 15-size scalpel (Swann Morton, Owlerton Green Sheffield, UK). The prospective craniotomy location above the primary somatosensory cortex (AP: −0.9 mm and ML: +3.0 mm relative to bregma) was marked with a biopsy punch (diameter 3 mm, Integra LifeSciences, Princeton, New Jersey, USA). The exposed skull around the area of interest was covered with a thin layer of dental acrylic (iBond Self Etch, Hereaus Kulzer, Hanau, Germany) and hardened with a LED polymerization lamp (Demi Plus, Kerr, Orange, CA, USA). A dental drill (Schick Technikmaster C1, Pluradent Frankfurt am Main, Germany) was used to thin the skull around the marked area. After applying a drop of sterile phosphate buffered saline (DPBS, Gibco, Life Technologies, Carlsbad, CA, USA) on the craniotomy the detached circular bone flap was removed using forceps (S&T Vessel Dilating Forceps - Angled 45°, FST, Heidelberg, Germany). Subsequently, SuperGrip forceps were used to remove carefully the dura mater (S&T Forceps - SuperGrip Tips, FST, Heidelberg, Germany), and the brain was rinsed with saline. A circular coverslip (3 mm diameter, thickness #0, VWR International, Radnor, PA, USA) was placed onto the craniotomy and glued to the skull with histoacrylic adhesive (Aesculap AG, Tuttlingen, Germany), applied via dental microbrushes (0.5 mm tip diameter, Microbrush International, Grafton, WI, USA). The exposed skull was covered with dental acrylic (Tetric Evoflow A1 Fill, Ivoclar Vivadent, Schaan, Liechtenstein) and a custom-made head-post was attached parallel to the window for head-fixing mice in subsequent imaging sessions. After surgery, the anesthesia was antagonized with a combination of Atipamezol (2.5 mg/kg), Flumazenil (0.5 mg/kg) and Naloxon (1.2 mg/kg) i.p.. Finally, mice received a subcutaneous dose of the analgesic Carprophen (7.5 mg/kg body weight, Rimadyl, Pfizer, New York, NY, USA) and were allowed to recover from surgery in a heating chamber.

### 6. Intravital 2-photon imaging

2-photon images were scanned as previously described^38^. Briefly, the anesthetized mouse was placed under an upright Zeiss LSM710 microscope equipped with a fs-laser (Mai Tai DeepSee, Spectra-Physics, Stahnsdorf, Germany), a 20x water immersion objective (W Plan-Apochromat 20x/1.0 NA, Zeiss) and a motorized stage. Ti:Sa laser (Chameleon Vision II) from 917 Coherent (Glasgow, Scotland) with an excitation wavelength of 830 nm and power of 5-22 % was used for detection. GAASP detector with an SP < 485 nm filter with master gain 600 for the donor channel and LP > 570 nm for the LNDs acceptor or FRET channel with master gain 600 were used. Images were taken as z-stacks (150 μm, 1024 × 1024, 8 bit, objective: W Plan-Apochromat 20x/1.0 DIC D=0.17 M27 75mm).

### 7. FRET in vivo ratiometric imaging

There were three types of LNDs which were used at FRET in vivo experiment: loaded with donor (F888), loaded with acceptor (Cy3) and with both donor and acceptor (FRET). All solutions were injected in volume 7.5 µl/kg body weight into the mice with implanted cranial windows. Mouse under MMF anesthesia were placed on the 2-PM stage. The LNDs were excited at 830 nm (and 740 nm for LNDs loaded with acceptor) and the emission was collected with SP < 485 nm (donor channel) and LP > 570 (acceptor/FRET channel) filters by non-descanned detectors (photomultiplier tube GaAsP, Zeiss). Three-dimensional z-stacks of 150 µm depth with 1 µm axial resolution and 1024×1024 pixels per image frame (0.6 µm/pixel) were acquired in multiphoton mode of the microscope over a time course of 120 min. The intensity was calculated at each channel as a total intensity of maximum intensity projection image using Fiji software. The FRET ratio was calculated using formula: FRETratio=FRET intensity/Donor intensity. Ratiometric analysis of LNDs integrity was performed in blood circulation 30-nm vs. 80-nm. Statistical analysis was performed using two-way ANOVA with Tukey correction for multiple comparison (n=3 for each group) with GraphPad® Prism (GraphPad Software, San Diego, CA, USA).

### 8. Nano-stroke, a transient mini-stroke mouse model

Stroke induction started one day after cranial window implantation. Throughout the experimental procedure, mice were anaesthetized with an intra-peritoneal injection of medetomidine (0.5 mg/kg), fentanyl (0.05 mg/kg), and midazolam (5 mg/kg) (MMF) (140/10 mg/kg) and placed on a heating pad to keep body temperature at 37 °C (Harvard Apparatus, Holliston, MA, USA). The head-holder of the mouse was fixed to a custom-made headpost and a neodymium-iron-boron (NdFeB) magnet was placed on the cranial window overlying the distal branches middle cerebral artery (MCA; Fig. 2D, i, ii). NdFeB magnets were custom-made (Zhenli Co. Ltd, Jiaozuo, Henan, China) with the following specifications: Cylindrical NdFeB magnets, grade N52, 1 mm long, 1.5 mm in diameter, magnetized along the cylindrical axis. Before injection, all solutions with nanoparticles were mixed thoroughly using a vortex shaker (VTX-3000L, LMS). Magnetic nanoparticles, Nanomag®-D (Micromod, Rostok, Germany), diameter 180 nm, PEG 2000 coated, were injected slowly at a volume of 6 µl/g body weight via a femoral artery catheter. Since iron oxide is a magnetite and each particle acts as an individual magnetic domain^39^, we expected to block the vessels via interaction between a magnet and blood-circulated MNPs. Safety, biocompatibility, and physical parameters of these MNPs were described previously^12^. Subsequently, injected MNPs (Fig. 2D, ii) into the blood stream were trapped in the magnetic field applied to the cerebral cortex, thereby occluding the underlying vessels as evidenced by the black colour filling the vessel lumen (Fig. 2D, iii). Reperfusion was then induced by removing the magnet. Ten minutes thereafter, the blood flow in the distal branches of middle cerebral artery (black arrow) as well as in most pial veins was restored (Fig. 2D, iv). Vessel occlusion was monitored and recorded under a stereo microscope M80 (Leica, Wetzlar, Germany), equipped with a video camera DFC 290 HP (Leica). After 1 hour the magnet was removed to induce reperfusion and a suspension of ultrabright FRET LNDs (7.5 µl/kg body weight) was injected. Systemic injection of fluorescent LNDs, was performed through same catheter used for MNPs. The platform used for performing the nano-stroke featured a custom heating pad and head-holder, which were compatible with the 2-PM stage. This allowed for easy relocation of the mouse to the microscope, enabling immediate intravital imaging of the reperfused area. Subsequently, mice were transferred under LSM 7 MP microscope (Zeiss, Oberkochen, Germany). LNDs were excited with 830 nm wavelength, laser power 5-11%, filters, as was described above, were SP < 485 nm and LP > 570 nm for donor and FRET respectively. Three-dimensional z-stacks of 150 µm depth with 1 µm axial resolution and 1024×1024 pixels per image frame (0.6 µm/pixel) were acquired in multiphoton mode of the microscope every 20 min over a time course of 60 min. Throughout the imaging session laser power was kept below 50 mW to avoid phototoxicity. Sham-operated animals received MNPs 20 min before imaging without magnet application. The microthrombi were identified visually as occluded vessels (lower-to-dark fluorescent intensity) or accumulated LNDs (higher fluorescent intensity).

### 9. Correlated light and ATUM Volume Scanning Electron Microscopy (CLEM)

In order to provide exact relocation of region of interest (ROI) from 2-PM to EM, the custom-made head post for head-fixation was used (Figure S05, A, white arrow), as described^40^. This enabled to relocate the imaging orientation from 2-PM to EM. Immediately after 2-photon imaging, mice were transcardially perfused with a mixture of 4% formaldehyde and 2.5% glutaraldehyde (Electron Microscopy Sciences, EMS) in 0.1 M sodium cacodylate buffer, pH 7.4. After 1 h perfusion mice were decapitated and the right parietal bone with the ipsilateral window was removed. Subsequently, the mouse head was attached to the stage of a vibratome (VT1000S, Leica Biosystems) via the head post to cut the brain tissue with thickness 80 µm parallel to the imaging plane (Figure S05A). This way, a single brain slice containing all cortical layers and the complete somato-sensory cortex was obtained. The cortical brain slice was incubated in the same fixative overnight at 4 °C, washed with PBS at the next day, cut at 50 µm thickness on a vibratome and subsequently stored in 0.1 M sodium cacodylate buffer at 4°C until the start of the postembedding. Coarse lateral relocation of the cortical ROI was guided by magnetic nanoparticles inspected under a standard binocular (Kern) and dissected at roughly 2×2 mm. A standard rOTO *en bloc* staining protocol^41^ was applied including postfixation in 2% osmium tetroxide (EMS), 1.5% potassium ferricyanide (Sigma) in 0.1 M sodium cacodylate (Science Services) buffer (pH 7.4). Staining was enhanced by reaction with 1% thiocarbohydrazide (Sigma) for 45 min at 40°C. The tissue was washed and incubated in 2% aqueous osmium tetroxide, washed and further contrasted by overnight incubation in 1% aqueous uranyl acetate at 4°C and for 2h at 50°C. Samples were dehydrated in an ascending ethanol series and infiltrated with epon-araldite resin (LX112, LADD).

In order to preserve the same imaging orientation for EM from the 2-photon experiment, the block was trimmed parallel to the cortical surface. The advantage of MNPs is to be seen label-free in light, fluorescent and electron microscopies^39^. Thereby, the correlation strategy relied on electron-dense magnetic particles and further anatomical landmarks like vascular patterns (Figure S05, B) and regions of MNP leftovers in the reperfused area, far outside of the ROI. The area of interest was exposed by trimming using TRIM90 (Diatome) on the ATUMtome (Powertome, RMC). In total, 1500 serial sections were collected starting from the pial surface with a 35° ultra-maxi diamond knife (Diatome) at a nominal cutting thickness and resulting axial resolution of 100 nm and collected on freshly plasma-treated (custom-built, based on Pelco easiGlow, adopted from M. Terasaki, U. Connecticut, CT), carbon nanotube (CNT) tape (Science Services). CNT tape stripes were assembled onto adhesive carbon tape (Science Services) attached to 4-inch silicon wafers (Siegert Wafer) and grounded by adhesive carbon tape strips (Science Services). EM micrographs were acquired on a Crossbeam Gemini 340 SEM (Zeiss) with a four-quadrant backscatter detector at 8 kV. In ATLAS5 Array Tomography (Fibics), wafer overview images were generated (2000 nm/pixel). In order to relocate the region imaged by light microscopy we acquired a data set covering 10 wafers (∼200 sections, covering 20 µm depth of tissue each) at a resolution of 0.2×0.2×4 µm. This coarse EM map was aligned (TrakEM2)^42^ and different cell types segmented (VAST)^43^. Building on this cellular map regions of interest surrounded but devoid of magnetic particle accumulations were selected for high resolution imaging at 20×20×100 nm voxel size. High resolution stacks were aligned, segmented and rendered (Blender)^44^. Accordingly, the first step was to identify the 2-PM field-of-view on a macro photograph of the cranial window (Figure S05, C, white arrows). This ROI was identified in low resolution EM using MNPs as fiducials and vascular patterns as endogenous landmarks (Figure S05, C, black arrows). Registration of the 2-PM and the low resolution SEM image volumes allowed us to target time-lapse features of interest like microthrombi formation or extravasation events (Figure S05, D, blue arrows) and acquire high-resolution volume SEM data thereof.

## Notes

### Competing Interest Statement

The authors have declared no competing interest.

### Summary of Updates

The title has been changed and some data added as well as transferred from supplementary to the main figures

