## Supplementary material for "Nanocarrier Drug Release and Blood-Brain Barrier Penetration at Post-Stroke Microthrombi Monitored by Real-Time Förster Resonance Energy Transfer-Based Detection System (FedEcs)": Supplemetary file

Title

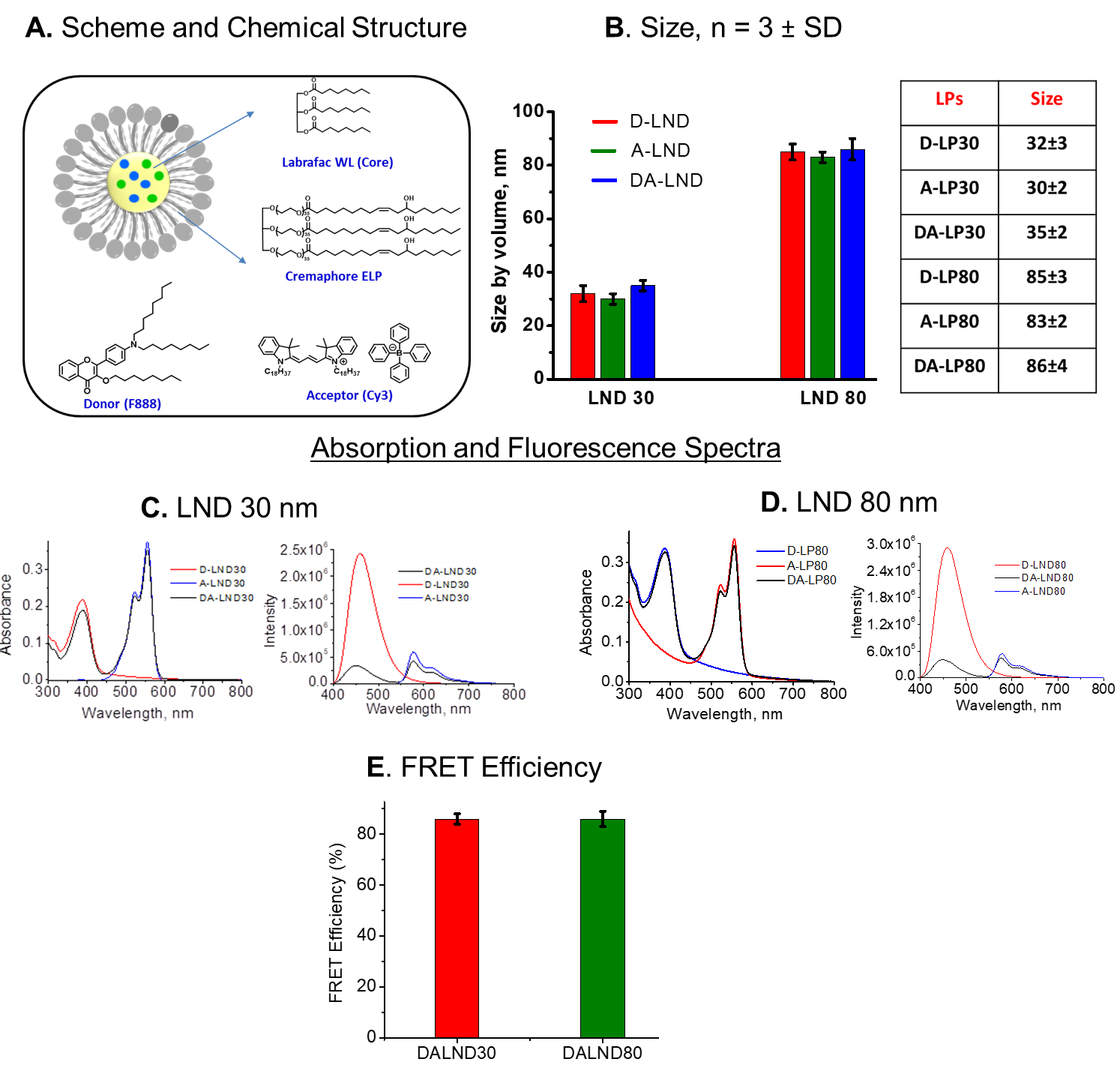
**Figure S1. Characteristics of lipid nanodroplets (LNDs) loaded with dyes to cause Forster resonance energy transfer (FRET). A.** Schematic representation of LNDs and chemical formulae for its components: Labrafac WL - core, Cremophore ELP - surfactant, F888 – donor dye, Cy3 - acceptor.  **B.** Size of LNDs measured by dynamic light scattering**.** D-LND – loaded solely by donor; A-LND – by acceptor; DA-LND – by pair of donor and acceptor. Absorption and fluorescence spectra of **C**) 30- and **D**) 80-nm LNDs (D alone, A alone and DA). **E.** FRET efficiency calculated from the emission spectra of donor in presence and absence of acceptor dyes.

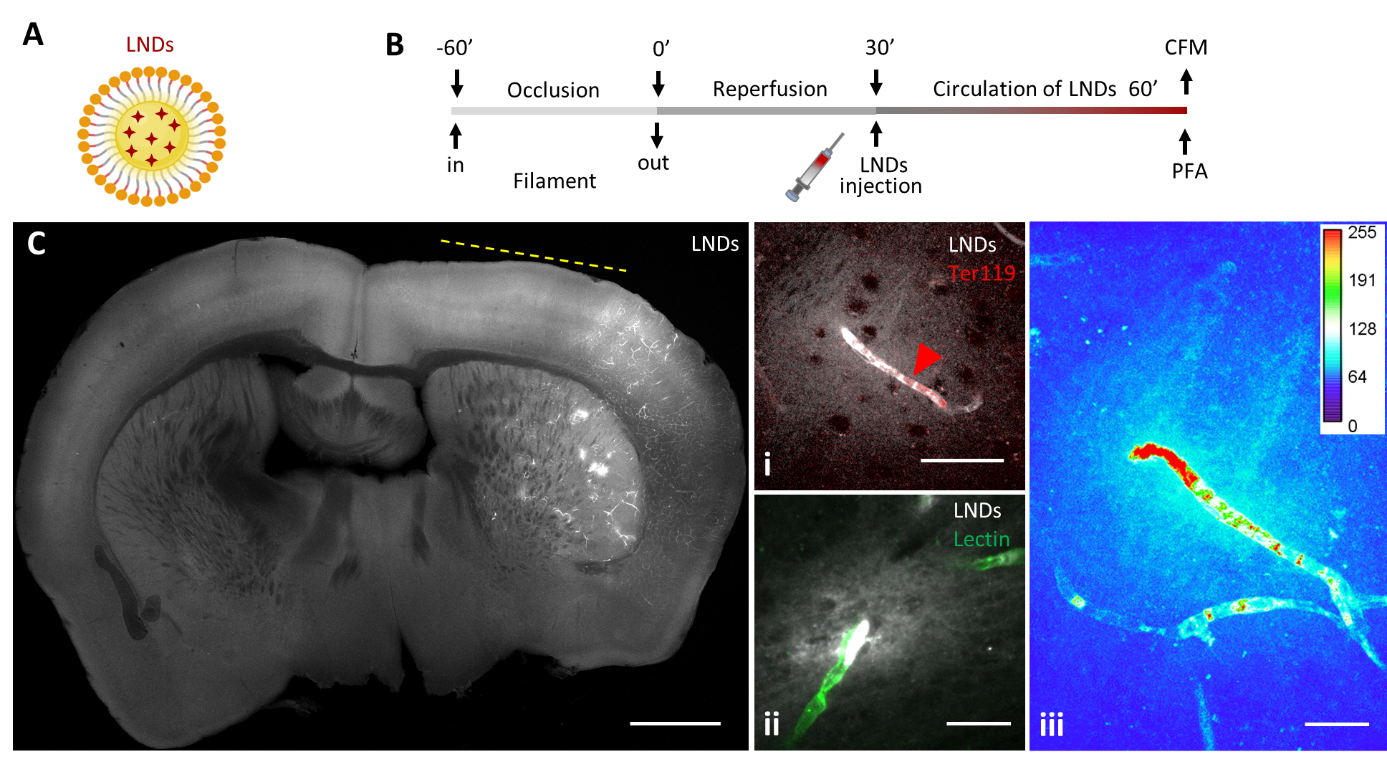
**Figure S2.** **Lipid nanodroplets accumulate in microvascular clots acutely after ischemic reperrfusion. A.** Schematic drawing of 30-nm lipid nano-droplets (LNDs) consist of an oil core (Labrafac WL) surrounded by surfactant (CreKolliphor ELP®) and loaded (red 4-pointed stars) with the fluorescent dye rhodamine bound to a hydrophobic counter-ion (R18/TPB). **B.** Experimental design. Injection of LNDs was performed within the acute reperfusion period after inducing stroke by the filament middle cerebral artery occlusion (fMCAo) mouse model. **C.** A representative confocal scan of a coronal section of the mouse brain with fMCAo, 60 minutes post-injection of 30-nm LNDs, scale bar 1mm. i – LNDs are trapped in an occluded vessel inside the re-perfused area, which is co-localized with erythrocyte (Ter119) stalls (red arrow), scale bar – 50 µm. ii - Extravasation of LNDs originates from the site of the blood clot. LNDs appear in the brain tissue beyond the vessels (Lectin) border. Scale bar 20 µm. iii - color-coding of a representative maximum intensity projection image, showing the grey values of accumulated and extravasated LNDs. Scale bar – 20 µm.

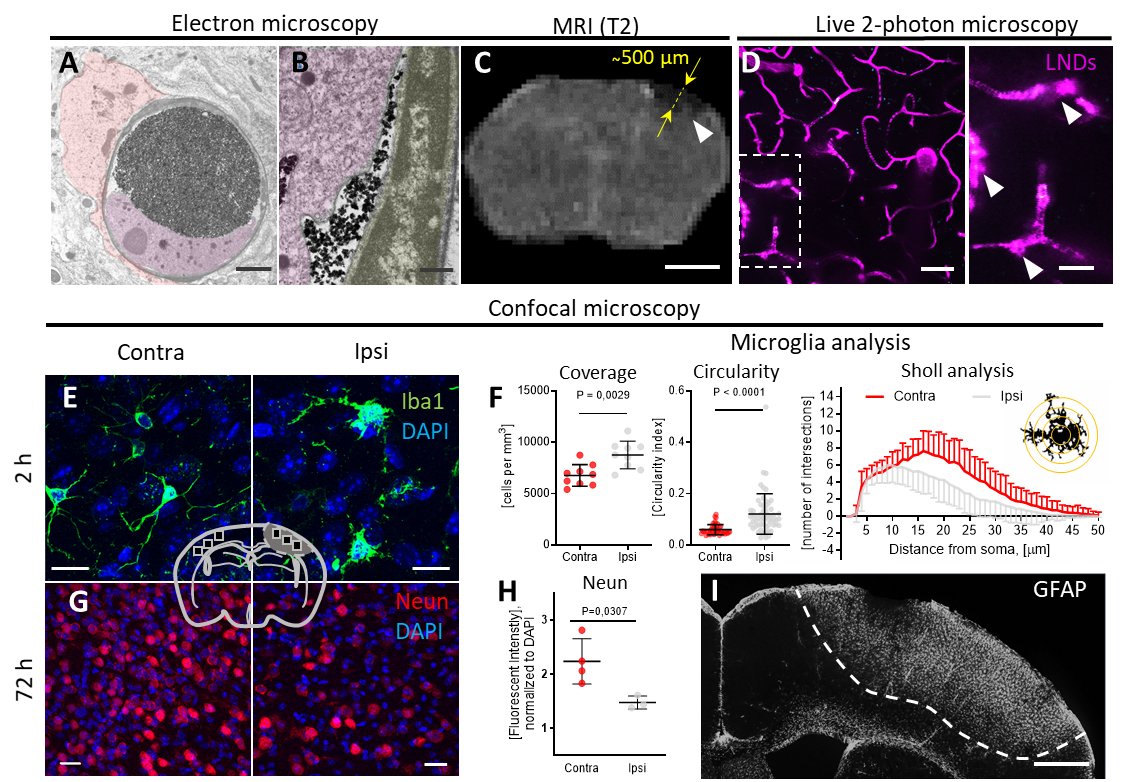
**Figure S3. Nano-stroke, as a new model for intravital imaging of ischemic brain.** **A.** Transmission electron microscopy (TEM) of a brain capillary occluded by MNPs (black dots) and macrophage (magenta). The occlusion is accompanied by astrocyte swelling (orange). Scale bar: 200 µm. **B.** TEM of occluded vessel shows that MNPs (black) do not destroy the endothelial cell luminal membrane (grey), but can be phagocytosed by macrophages (magenta). Scale bar: 500 nm. **C.** T2-weighted magnetic resonance imaging (MRI) demonstrates accumulation of MNPs deep in cortical tissue after one hour of ischemia. White arrow – accumulated MNPs. Yellow arrows- depth of the ischemic area within the cerebral cortex (500 µm). Scale bar – 2 mm. **D.** 2-photon intravital microscopy of LNDs circulated in reperfused cerebral microvessels after removal of the micro magnet. Maximum intensity projection. Dashed line – zoomed area; white arrows - clots with accumulated LNDs. Scale bars: 50 µm, zoomed area: 20 µm. **E.** Confocal images of a coronal section of mouse brain stained for microglia (Iba1) in the area under the magnet (Ipsi) and homotopical area (Contra) 2 hours after reperfusion. **F.** Coverage, fractal (circularity) and sholl analyses of microglia. The coverage is higher in the ipsilesional area compared to contralesional. Data are presented as mean ± standard deviation (SD), n=9 regions of interest (ROI). Student t-test was used. The fractal analysis revealed that microglia in ipsilateral area show a higher circularity index, indicating a shift from resting to activated microglia. Data are presented as mean ± SD, n=45-58 cells. Student t-test was used. The Sholl analysis also shows decreased ramification, i. e. more active cells, in the ipsilateral part. Data are presented as mean ± SD; n=45-58 cells. Two-way ANOVA with Tukey correction for multiple comparison was used. **G.** Confocal images of a coronal section of mouse brain stained for neurons (Neun) 72 hours post-reperfusion **H** Quantification of Neun fluorescent intensity in the ROI of ipsilateral part compared to contralateral shows the neuronal degradation in the area underneath the magnet. **I.** A representative confocal image of a whole lesion in coronal section of mouse brain stained for glial fibrillary acidic protein (GFAP) showing gliosis. Scale bar: 0.5 mm.

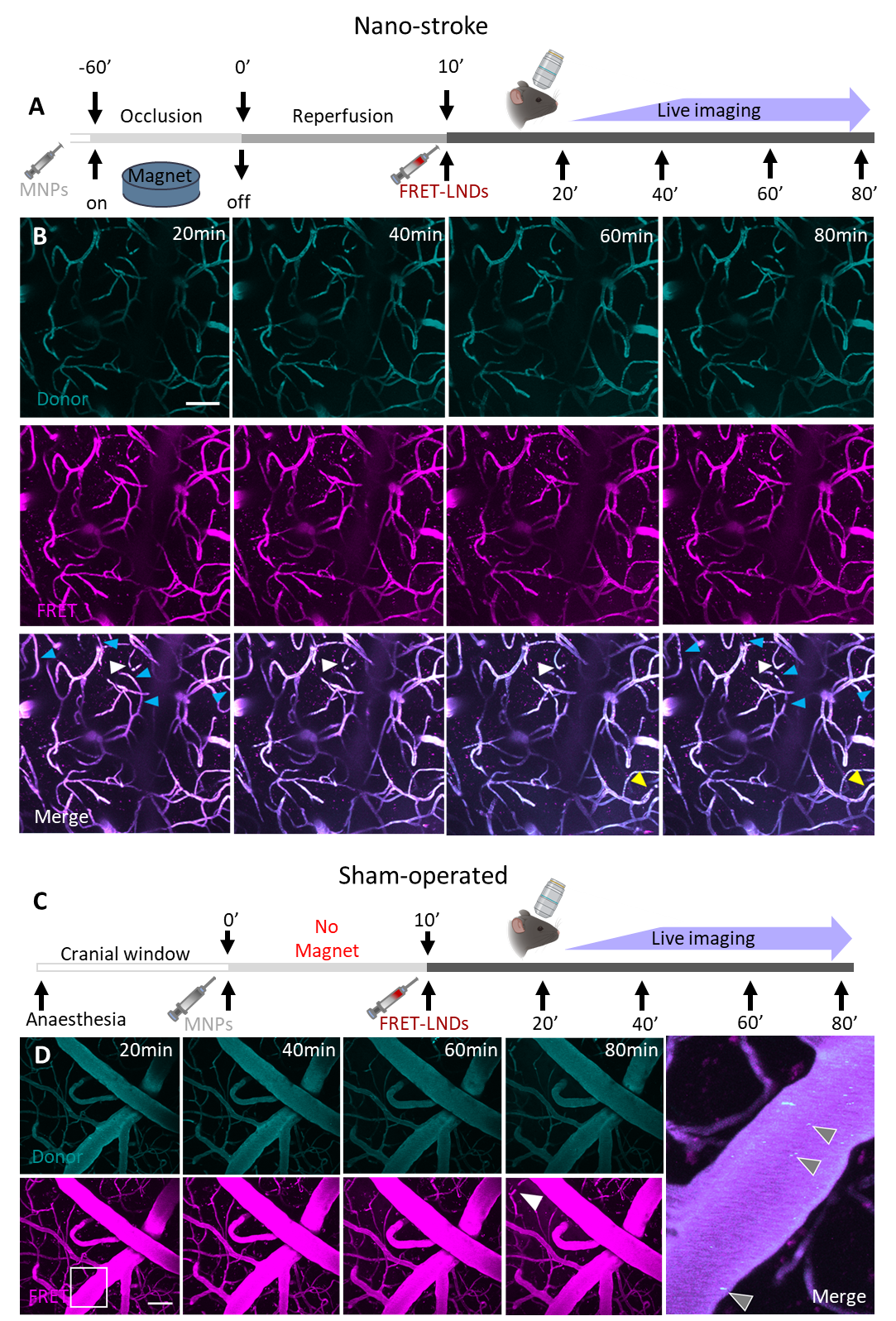

**Figure S4. Spontaneous microvascular clot formation in nano-stroke model *vs.* sham. A.** Experiment design as described at Fig. 2A with additional longitudinal intravital imaging during 60 minutes. **B.** A representative maximum intensity projection (MIP) of Z-stack of longitudinal intravital imaging of cortical area after nano-stroke model in mouse injected with FRET lipid nanodroplets (LNDs) as shown at Fig. 2A. Blue – donor channel; magenta – acceptor channel. White arrows – permanent clots; blue arrows – newly formed spontaneous clots; yellow arrows – self-resolving clots. **C.** Experiment design for sham-operated mouse. Mice went through the same procedure as nano-stroked animals, but without occlusion period by a magnet on the top of the glass cranial window. **D.** A representative MIP of Z-stack of longitudinal intravital imaging of cortical area of sham-operated mouse injected with FRET LNDs. Blue – donor channel; magenta – acceptor channel. White arrow - single vascular occlusion. Grey arrows at the zoomed and merged area – circulated magnetic nanoparticles. Scale bars - 50 µm.

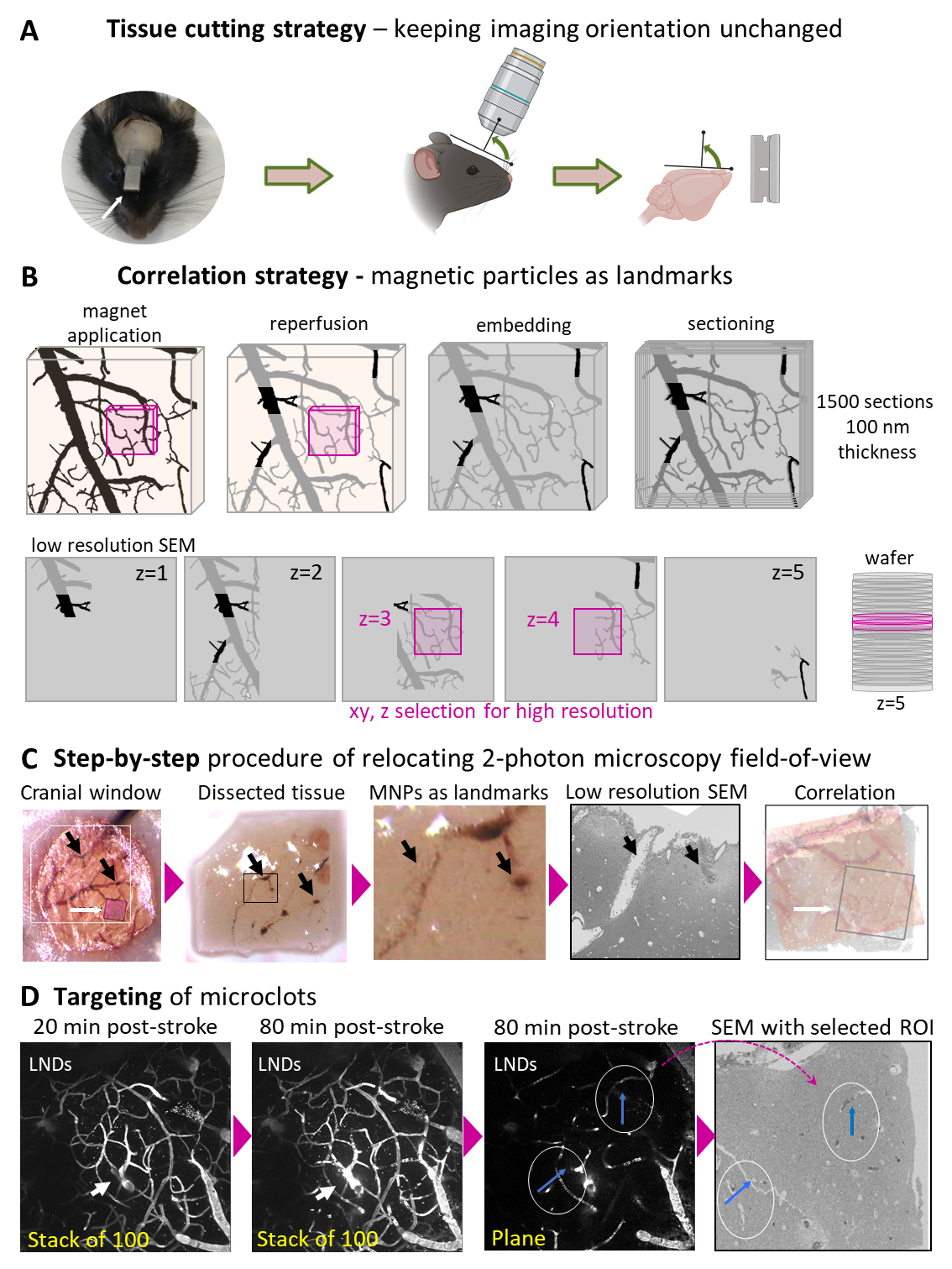

**Figure S5. Strategy for correlated 2-photon (2-PM) and ATUM volume SEM based on magnetic nanoparticle fiducials. A.** Cutting strategy ensuring the conservation of the imaging orientation from 2-PM to SEM and thereby facilitating correlation and relocation. Custom made head holder compatible with 2-PM station and vibratome stage (white arrow). **B.** Schematics of the correlation approach using MNPs. Top row: A cube of tissue within the cranial window is shown. After magnet application vessels are filled with MNPs (black) while most of the MNPs are removed after reperfusion (grey). After embedding for EM, MNPs are visible in EM micrographs. The actual region of interest (pink) was determined at a site without MNPs but with surrounding MNPs serving as fiducials. Bottom row: Serial sections are collected, mounted on wafers (right) and imaged at low resolution in the SEM. The vascular pattern has to be reconstructed in order to match it to the initial 2-PM image. The right location in z (pink) is found through low resolution wafer screening. **C-D**. Step-by-step procedure for ROI relocation of a microvascular clot. **C.** Finding the field-of-view. From left to right: light microscopy image of the cranial window, binocular image of the dissected tissue and a high magnification view thereof; low resolution SEM micrograph of the same region showing occluded (blue) and empty (black) large vessels; correlation of LM with SEM (boxed region is shown below). **D.** Relocating the microclots. The clot was not present at 20 min post-stroke but emerged 60 min later as observed by 2-PM (white arrow). From left to right: summed 2-PM images at 20 min, 80 min and single plane. Right: SEM overview image of the same region. Occluded vessels (blue arrow). **D**. MNP – magnetic nanoparticles, Nanomag-D®; SEM – scanning EM; LND – lipid nanodroplets, 30 nm.

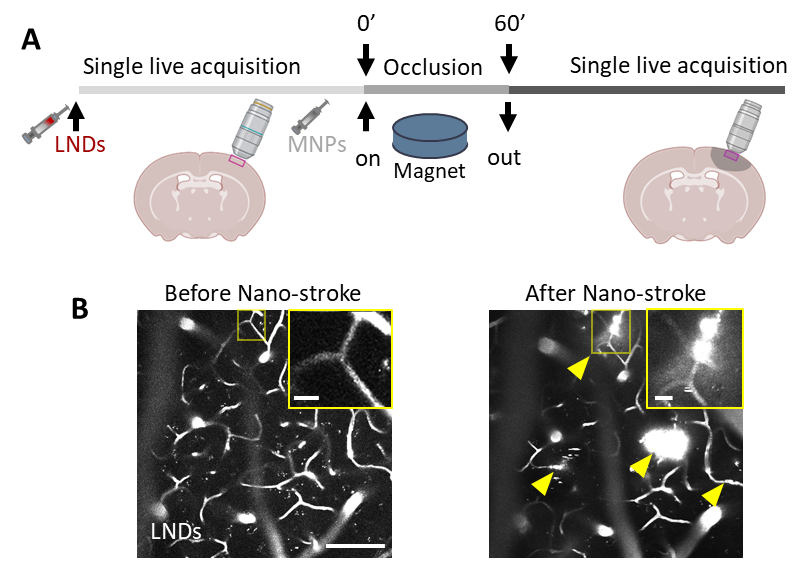

**Figure S6. Mechanical vessel occlusion by magnetic nanoparticles (MNPs) causes extravasation of lipid nanodroplets (LNDs). A.** Experiment design: LNDs were injected into a healthy mouse with an implanted cranial window, followed by a single imaging session using 2-photon microscopy to observe the LNDs circulating in the brain's vascular system. Then Nano-stroke during 60 min was applied and mouse immediately was placed under 2-photon microscope to perform live imaging of same region. **B.** Proof-of-principle that LNDs-based brain-targeted drug-delivery is possible through artificial vascular occlusions. Insert – zoomed area outlined by yellow line. Before occlusion – LNDs were circulating inside intact cerebral vessels; after occlusion LNDs extravasated into the parenchyma immediately after the removal of the neodymium mini-magnet. Yellow arrows – spots of extravasation. Scale bar – 100 µm, insert – 10 µm.

**Movie S01.** Comparison of representative Z-stacks of correlated microthrombi (Fig. 3B, D) of interest acquired using intravital 2-photon microscopy (left) and scanning electron microscopy (right).

**Movie S02.** 3D model of the correlated microthrombi, which were formed in different time points. The model was reconstructed from the volume SEM data set as shown in Fig. 3B-F. LNDs (green), endothelium (transparent grey), pericyte (yellow), astrocytes (sky blue), microglia (dark blue), platelet (magenta), erythrocytes (red).

**Movie S3.** 3D model of the correlated extravasation spot associated with microthrombus. The model was reconstructed from the volume SEM data set as shown in Fig. 3G-H. LNDs (dark blue), endothelium (transparent grey), fibrin (dark grey), pericyte (green), microglia (sky blue), immune cell (yellow), magenta (platelets), erythrocytes (red).
